# Differential effects of aging on regional corpus callosum microstructure and the modifying influence of pulse pressure

**DOI:** 10.1101/2023.10.31.562251

**Authors:** Jessica N Kraft, Stephanie Matijevic, David A Hoagey, Kristen M Kennedy, Karen Rodrigue

## Abstract

The corpus callosum is composed of several subregions, distinct in cellular and functional organization. This organization scheme may render these subregions differentially vulnerable to the aging process. Callosal integrity may be further compromised by vascular risk factors, which negatively influence white matter health. Here, we test for heterochronicity of aging, hypothesizing an anterior-to-posterior gradient of vulnerability to aging that may be altered by effects of vascular health. In 174 healthy adults across the adult lifespan (mean age=53.56 ± 18.90, range=20-94 years old, 58.62% women), measures of pulse pressure and diffusion-weighted imaging were conducted. A deterministic tractography approach was utilized to extract fractional anisotropy (FA), radial diffusivity (RD) and axial diffusivity (AD) from each of five callosal subregions, serving as estimates of microstructural health. General linear models tested effects of age, hypertension, and pulse pressure on these metrics. We observed no significant effect of hypertensive diagnosis on callosal microstructure. We found a significant main effect of age and an age x pulse pressure interaction whereby older age and elevated pulse pressure were associated with poorer FA, AD, and RD. Age effects revealed non-linear components and occurred along an anterior-posterior gradient of severity in the callosum. This gradient disappeared when pulse pressure was considered. These results indicate that age-related deterioration across the callosum is regionally variable, and that pulse pressure, a proxy of arterial stiffness, alters this aging pattern.

**Significance Statement:** Aging is associated with microstructural changes in the corpus callosum, the largest white matter tract in the brain. Additionally, vascular factors, such as hypertension and pulse pressure, affect corpus callosum microstructure. However, it is unclear whether these factors uniformly impact the corpus callosum throughout aging. The current study aimed to characterize patterns of aging and assess the role of hypertension and pulse pressure across different subregions of the corpus callosum. We found an age-related gradient on corpus callosum microstructure, with the most pronounced impact on anterior regions. However, this gradient was not found when pulse pressure was considered. These findings suggest that subregions are differentially sensitive to age-related decline, and pulse pressure modifies and exacerbates these declines.

## Introduction

The corpus callosum (CC) subregions are characterized by distinct axonal diameters, fiber densities, and myelination levels (Aboitiz et al., 1992). Projections from sets of homologous cortical areas traverse the callosal subregions (Hofer & Frahm, 2006; Huang et al. 2005), enabling each subregion to facilitate distinct aspects of cognitive operations (Baird et al., 2005). Specific subregions reach maturation at different points in development (Lebel et al., 2010) and each callosal subregion may be differentially vulnerable to aging (Michielse et al., 2010; Sullivan, Adalsteinsson, & Pfefferbaum 2006).

Magnetic resonance imaging (MRI) allows for indirect assessments of white matter microstructural health along the corpus callosum via diffusion tensor imaging (DTI). DTI evaluates axonal attributes of white matter by extracting information on the orientation and magnitude of water diffusion from diffusion weighted-MR images (Basser & Pierpaoli, 1996). Fractional anisotropy (FA), a metric of the directionality of diffusion, varies across the callosum (Hasan et al., 2009; Ota et al., 2006), reflecting differences in axonal composition among subregions. Studies have demonstrated non-linear associations between FA values and age, characterized by increases in FA from birth to young adulthood and steady decreases through adulthood (Bendlin et al., 2010; Hsu et al. 2010). Age-related losses in FA in healthy adults may be greatest for anterior white matter regions and least severe for posterior brain regions (Bennet et al., 2013; Kennedy & Raz, 2009b). FA values in the CC specifically follow non-linear patterns of development and decline (Lebel et al., 2010; McLaughlin et al., 2007). This variability suggests an anterior-to-posterior deterioration gradient of aging (Head et al., 2004; Madden, Bennett & Song, 2009), likely due to a mirroring of development of these subregions of the CC, sometimes referred to as retrogenesis (Brickman et al., 2012; Stricker et al., 2009), or the last-in first-out principle based on the developmental heterochronicity of the CC subregions (Raz, 2000; Raz & Kennedy, 2009).

Additional DTI metrics, such as radial diffusivity (RD) and axial diffusivity (AD), are thought to indicate a more specific form of white matter damage than that captured in FA (Metwalli et al, 2010). Animal research suggests that increases in RD may represent de/dysmyelination, whereas increases in AD may represent Wallerian degeneration and total axonal loss (Song et al., 2003). Typically, non-pathological brain aging is associated with increases in RD and minimal-to-no increase in AD, though the literature is mixed (Bennet et al., 2010, Ingo et al., 2021, Bender & Raz, 2015).

Along with age, vascular health influences white matter integrity. Hypertension specifically is associated with a higher incidence of white matter hyperintensities and greater age-related reductions in white matter volume and FA within the genu of the CC (Burgman et al., 2010). Incremental elevations in blood pressure may exert negative effects on white matter microstructure in even normotensive populations (Kennedy & Raz, 2009; Maillard et al., 2012; Salat et al., 2012, Hoagey et al., 2021). Moreover, the effects of vascular risk factors may be regionally selective, though the nature of regional specificity is under debate. Burgman et al. (2010) reported stronger effect of hypertension in the genu and body of the callosum than in the splenium, whereas Gons et al. (2012) reported a stronger effect in the splenium. Greater baseline vascular risk (such as clinical diagnoses of hypertension or hypercholesterolemia) was associated with substantial longitudinal declines of FA in the body and splenium of the CC (Williams et al., 2019).

Most work examining the association of age and vascular health on white matter microstructure has focused on older adult cohorts with truncated age ranges. While older age and hypertensive risk factors have been associated with degradation of callosal microstructural integrity (Gons et al., 2012; Burgman et al., 2010; Williams et al., 2019), the impact of these risk factors over the lifespan remain unclear. Research has suggested this degradation of callosal microstructure occurs in a non-linear fashion throughout the aging process (Kennedy & Raz, 2009a, Bendlin et al., 2010). Given the prevalence of blood pressure elevations and arterial stiffness in the aging population, assessing susceptibility to microstructural damage throughout the lifespan is imperative. This is especially true in younger and middle-aged adults, for whom interventions to preserve callosal structure and vascular risk reduction may be most impactful. Therefore, the goal of the following study was to investigate the effects of age, hypertension, and pulse pressure -- a proxy for arterial stiffness -- on callosal microstructural integrity throughout the adult lifespan. Specifically, we sought to characterize patterns of aging across individual subregions of the CC and evaluate the effects of hypertension and pulse pressure to determine whether age and vascular health exerted regionally differential influence on callosal microstructure.

## Methods

### Participants

Participants included 174 healthy, right-handed, native English-speaking adults aged 20-94 (102 women, 72 men) who were recruited for the study through advertisements and flyers from the Dallas - Fort Worth metroplex and were compensated for their participation. Participants were screened to be free from cardiovascular disease, diabetes, cancer, neurological and psychiatric disorders, and drug and alcohol abuse through both phone and mail-in health questionnaires prior to entry into the study. Participants also underwent screenings for vision, hearing, and cognitive impairment at the first study visit. Those with a score > 16 on the Center for Epidemiological Studies Depression Scale (Radloff, 1977) or a score < 26 on the Mini Mental Status Exam (Folstein, Folstein, & McHugh, 1975) were considered ineligible. Written informed consent was obtained in accordance with the guidelines set by The University of Texas at Dallas and The University of Texas Southwestern Institutional Review Boards. Participants underwent two cognitive testing sessions and an MRI session. See Table 1 for participant demographic information.

**Table 1.** Sample Demographics and Blood Pressure Measurements.

| **Demographic characteristics** |  |
| --- | --- |
| Number of participants | 174 |
| Age, mean ± SD | 53.56 ± 18.90 |
| Sex, N (%) |  |
| Men | 72 (41.38%) |
| Women | 102 (58.62%) |
| Years of Education, mean ± SD | 15.50 ± 2.50 |
| MMSE, mean ± SD | 29.02 ± 0.85 |
| CESD, mean ± SD | 4.32 ± 3.79 |
| BMI, mean ± SD | 27.30 ± 5.13 |
| Smokers, N (%) | 7 (4.07%) |
| High Cholesterol, N (%) | 36 (20.93%) |
| Diagnosed Hypertensives, N (%) | 38 (21.84%) |
| Systolic BP, mean ± SD | 126.37 ± 16.54 |
| Diastolic BP, mean ± SD | 75.26 ± 9.54 |
| Heart Rate, mean ± SD | 68.43 ± 9.84 |
| Pulse Pressure, mean ± SD | 51.11 ± 12.06 |
| Mean Arterial BP, mean ± SD | 92.29 ± 10.93 |

*Note.* MMSE -- Mini Mental Status Exam; CESD -- Center for Epidemiological Studies Depression Scale; BMI -- Body mass index; BP -- Blood pressure.

### Blood Pressure Measurements

Participants provided information on hypertension diagnoses and antihypertensive medication use during the initial screening process. Systolic blood pressure (BP), diastolic BP and heart rate readings were collected during both cognitive testing sessions and the MRI session. Readings for all three visits were averaged and used to calculate mean pulse pressure (systolic minus diastolic), which is indicative of arterial stiffness in large arteries. Participants with a previous diagnosis of hypertension by a physician were classified as diagnosed hypertensives, while participants with either mean systolic BP ≥ 140 mmHg or mean diastolic BP ≥ 90 mmHg and no previous hypertension diagnosis were classified as undiagnosed hypertensives.

### Neuroimaging Acquisition and Pre-processing

All participants were scanned on the same 3T Philips Achieva MR scanner (Philips Medical Systems, Best, Netherlands) with a 32-channel head coil using SENSE encoding at The University of Texas Southwestern Medical Center Advanced Imaging Research Center. A 3D T1-weighted MP-RAGE image was acquired with a single turbo field echo sequence (160 sagittal slices, TR = 8.3 ms, TE = 3.8 ms, flip angle = 12°, T1 = 1100 ms, voxel size = 1 x 1 x 1 mm³, acquisition time = 3:57 minutes). Diffusion weighted (DW) scans were obtained using a single shot echo-planar imaging sequence (65 axial slices, 30 gradient directions, b = 1000, 1 non-diffusion weighted b_0_, TR = 5611 ms, TE = 51 ms, flip angle = 90°, voxel size = 2 x 2 x 2.2 mm³, slice thickness = 2.2 mm, acquisition time = 4:19 minutes).

T1 images were skull stripped (via BET; Smith, 2002), intensity bias corrected, and registered to Montreal Neurological Institute 1 mm template space (Montreal Neurological Institute, McGill University, Canada) via ANTS (Avants et al., 2011). Diffusion images were visually inspected for scanner artifacts and brain abnormalities. Automated software was used to detect motion, susceptibility, and eddy current distortions, while corrections were applied by either using a linear registration of each diffusion gradient to the non-diffusion weighted b_0_ or removing corrupted gradients from analysis via DTIPrep (Liu, 2010). Diffusion directions were adjusted to account for reorientation of individual gradients (Leemans & Jones, 2009). DTI scalar and tensor maps were calculated using the DSI Studio software program (Yeh, 2013).

### Callosal Segmentation and Fiber Tracking

Diffusion indices were extracted from callosal segments using an approach which elucidated white matter fibers via deterministic tractography within a region-of-interest (ROI) defined as the corpus callosum. Using previous parcellation schemes as a guide, the corpus callosum was manually traced on the 1 mm MNI template to create five separate topographic subregions of genu, anterior midbody, posterior midbody, isthmus, and splenium (cf Hofer and Frahm 2006; Witelson 1989). Non-linear registration algorithms placed each callosal division in participant’s native diffusion space to define the region with which to restrict tractography results (Avants et al., 2011). Interhemispheric projections were resolved via streamline deterministic tractography using the midsagittal plane as an ROI to restrict tracking (Yeh, 2013). Additionally, to remove extraneous fibers projecting into the cingulum, we isolated the cingulum bundle separately for use as an exclusion mask. Resulting fibers were divided into the five callosal subregions based upon their location within the manually traced ROIs. This approach was used to ensure that diffusion metrics would only be extracted from viable fibers which were both interhemispheric and within a subdivision of the corpus callosum. Mean fractional anisotropy (FA), radial diffusivity (RD), and axial diffusivity (AD) were extracted from each of the five callosal segments. See Figure 1 for illustration of the tractography-guided segmentation results.

**Figure 1.**
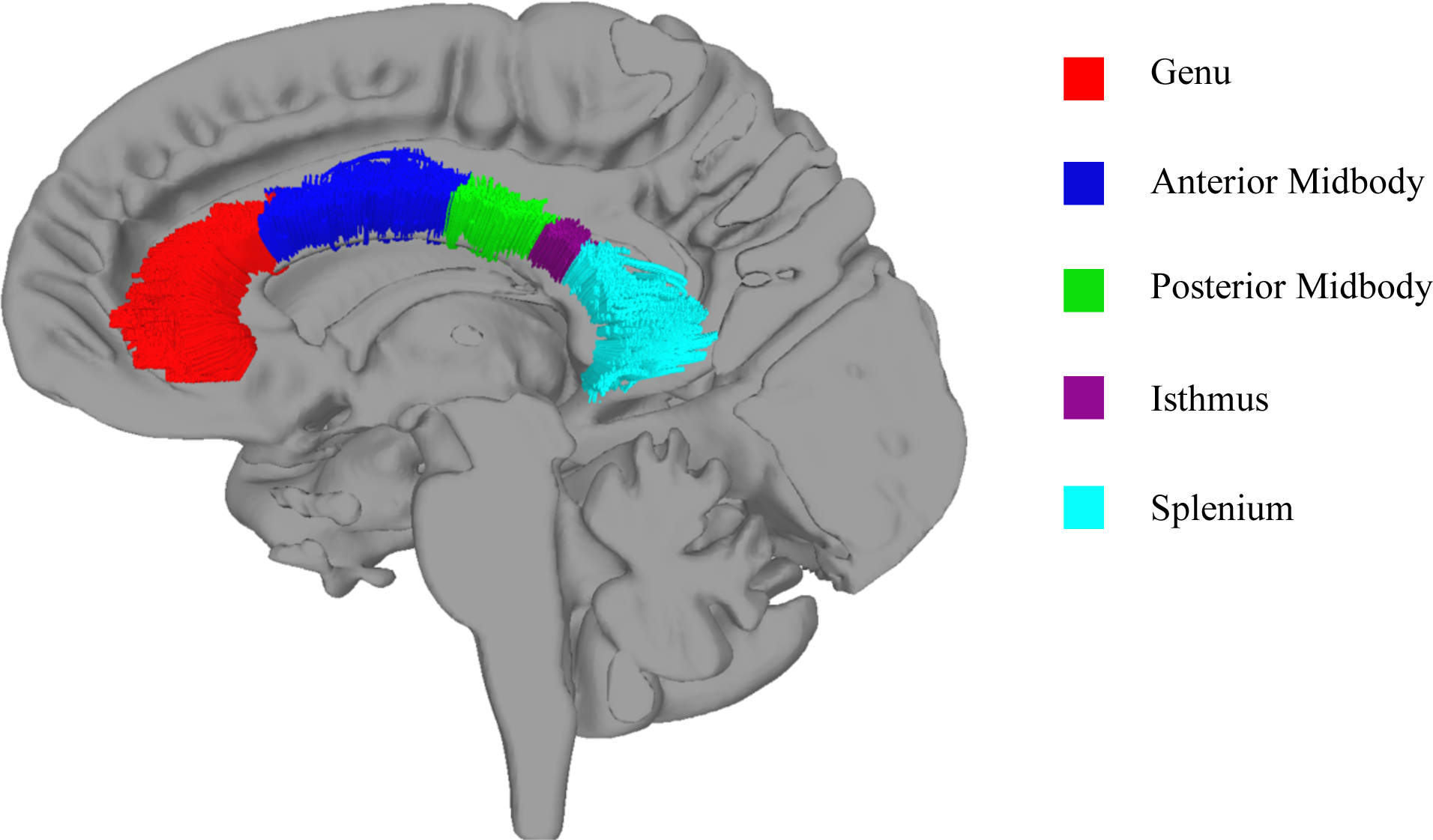
Example of callosal subregion ROI tracts registered to a subject’s native b_0_ image. Color represents each callosal subregion tract fibers.

## Results

Descriptive statistics for the sample are provided in Table 1. Hypertensives, both undiagnosed and diagnosed, did not differ from normotensives in years of education, MMSE or CESD scores, but did differ greatly in blood pressure levels, as would be expected.

A series of general linear models (GLMs) was performed to examine the effects of age, hypertension diagnosis, and pulse pressure on measures of white matter microstructural integrity extracted from the callosum subregions. In every model, age was set as a continuous variable, sex as a categorical variable, and mean values of FA, RD, or AD as dependent variables with ROI (subregions 1-5) as a within-subjects factor. Age and pulse pressure were centered at the mean. Sex was not found to have significant (*p* < 0.05) effects or interactions in any of the models and was removed along with other nonsignificant interactions from final models to conserve statistical power.

### Effects of Age on Regional CC Tract Diffusion Metrics

In the GLMs for age as the sole independent variable, we observed a significant main effect of age for FA (F(1,173)=78.50, p<0.0001), RD (F(1,172)=106.67, p<0.0001) and AD (F(1,173)=10.40, p=0.002). There was a significant age x ROI effect for FA (F(7,167)=15.71, p<0.0001), RD (F(7,166)=23.09, p<0.0001), and AD (F(7,167)=5.65, p<0.0001), indicating that age may have a differential effect on the subregions of the callosum. Of note, one individual was excluded from RD analyses for extreme outliers, defined as having a value of > 3 times the interquartile range (IQR) in each subsegment of the CC. One additional participant was excluded from AD analyses within the genu of the CC for an extreme outlier, defined as a value > 3 times the IQR.

To decompose these significant interaction effects, we produced zero-order correlations (Table 2) for the association between age and mean FA, RD, and AD values in each subregion. FA negatively correlated with age for all five subregions, while RD and AD positively correlated. All associations except for AD in the posterior midbody and splenium were significant. See Figures 2-4.

**Figure 2.**
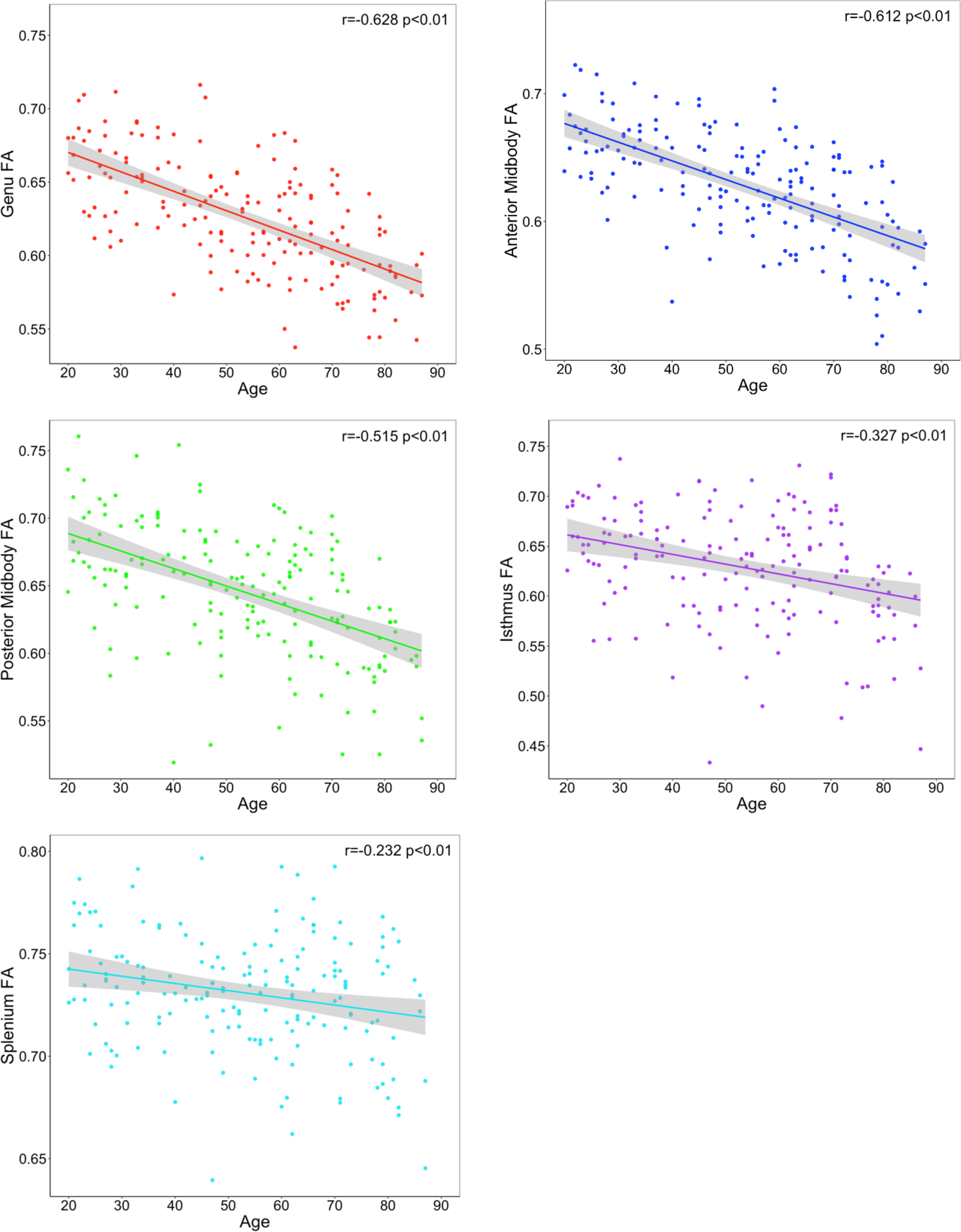
Effects of age on regional tract FA across the corpus callosum segments

**Figure 3.**
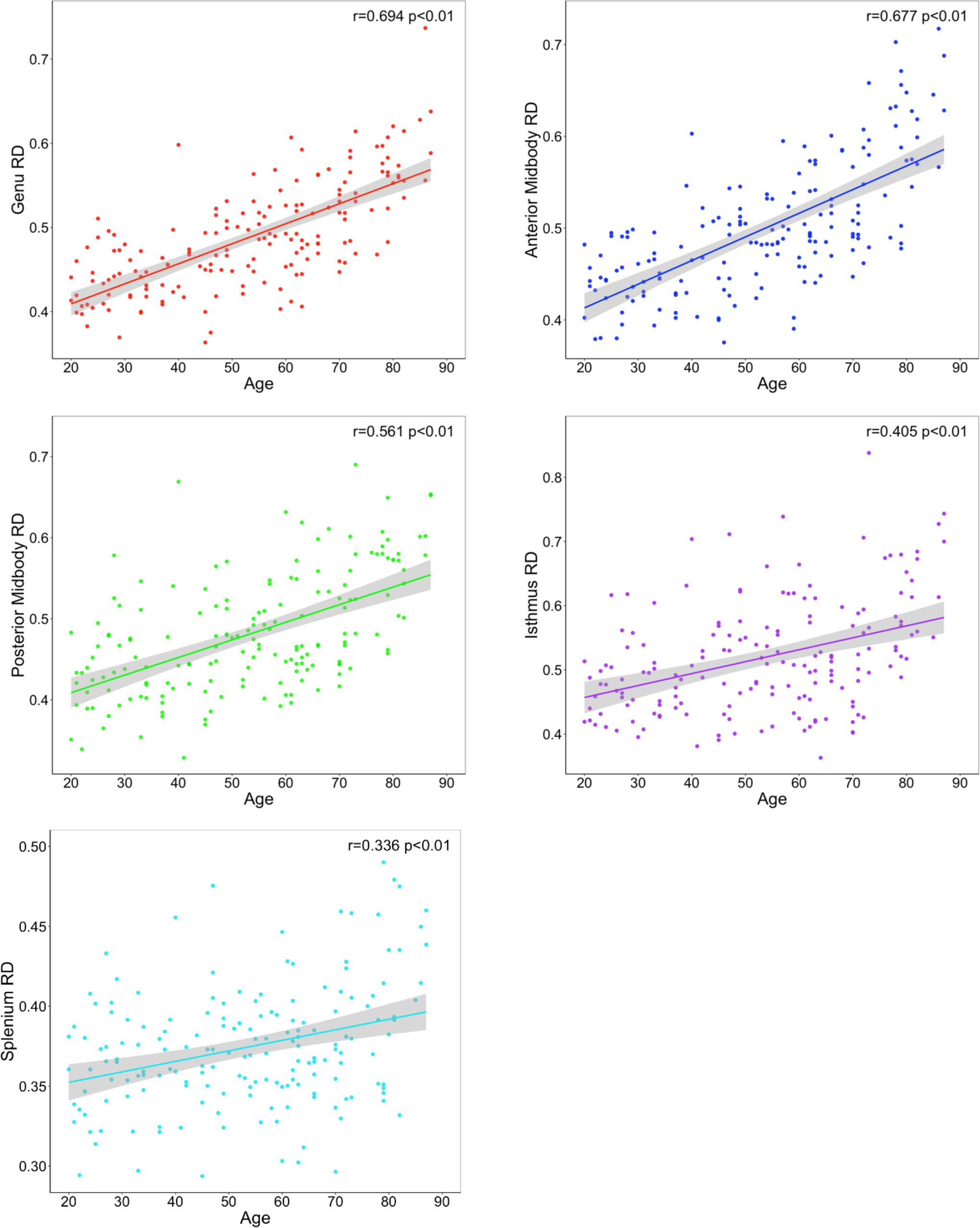
Effects of age on regional tract RD across corpus callosum segments

**Figure 4.**
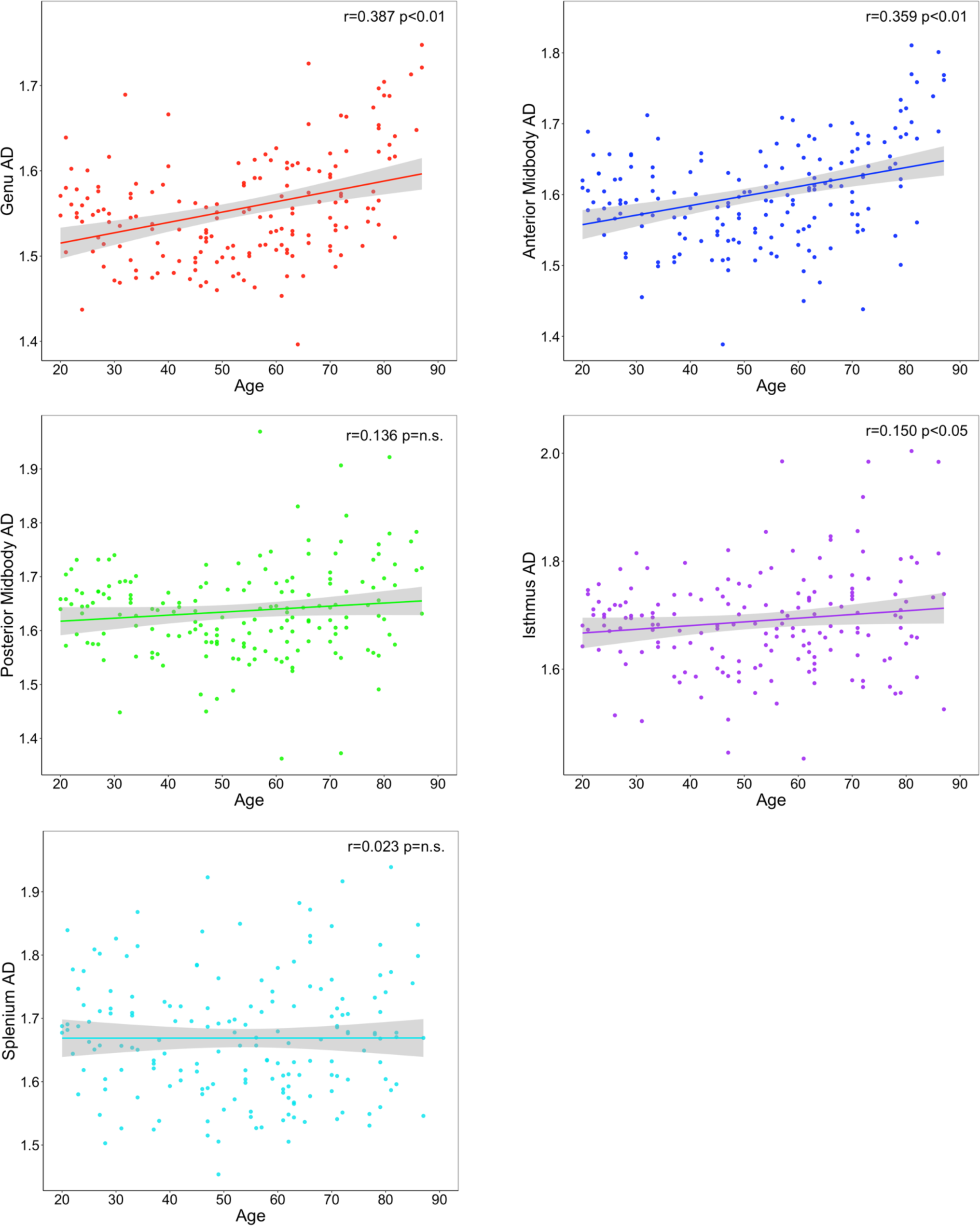
Effects of age on regional tract AD across corpus callosum segments

**Table 2.** Associations between age and DTI metrics within CC segment tracts.

|  | **FA** |  | **RD** |  | **AD** |  |
| --- | --- | --- | --- | --- | --- | --- |
| **Segment** | **Mean ± SD** | ***r*** | **Mean ± SD** | ***r*** | **Mean ± SD** | ***r*** |
| Genu | 0.63 ± 0.04 | -0.628* | 0.49 ± 0.06 | 0.694* | 1.56 ± 0.07 | 0.387* |
| Anterior Midbody | 0.63 ± 0.04 | -0.612* | 0.50 ± 0.07 | 0.677* | 1.60 ± 0.07 | 0.359* |
| Posterior Midbody | 0.65 ± 0.05 | -0.515* | 0.48 ± 0.07 | 0.561* | 1.64 ± 0.09 | 0.136 |
| Isthmus | 0.63 ± 0.06 | -0.327* | 0.52 ± 0.09 | 0.405* | 1.69 ± 0.09 | 0.150** |
| Splenium | 0.73 ± 0.03 | -0.232* | 0.38 ± 0.04 | 0.336* | 1.67 ± 0.10 | 0.023 |

*Note.* Pearson's correlations, uncorrected
\* $p < 0.01$
\*\* $p < 0.05$

We used Steiger’s *Z* test to compare the magnitude of these significant findings of age on FA, RD and AD between different subregions. Overall, for all metrics, correlation with age was strongest in the genu and weaker in more posterior subregions in a stepwise fashion across the callosum. This gradient appears to be the strongest for FA and RD. The circular barplot in Figure 5 illustrates the differential impact of age on regions of the CC for FA, RD, and AD values. For FA, there were statistically significant differences on the age and CC association between the genu and splenium, the genu and isthmus, the anterior midbody and splenium, the anterior midbody and isthmus, and the posterior midbody and splenium (all Steiger’s Z p-values <0.001), and the posterior midbody and isthmus (p<0.05). We observed a similar stepwise pattern with RD metrics: between the genu and isthmus, genu and splenium, anterior midbody and isthmus, anterior midbody and splenium, and the posterior midbody and splenium (all p-values <0.001), and between the genu and posterior midbody, and isthmus and splenium (all p-values<0.05). A similar pattern exists for AD, albeit less pronounced: between the genu and the splenium, the genu and the posterior midbody, and the anterior midbody and splenium (all p-values< 0.001), and between the genu and isthmus, anterior midbody and isthmus, and anterior midbody and splenium (all p-values <0.05).

**Figure 5.**
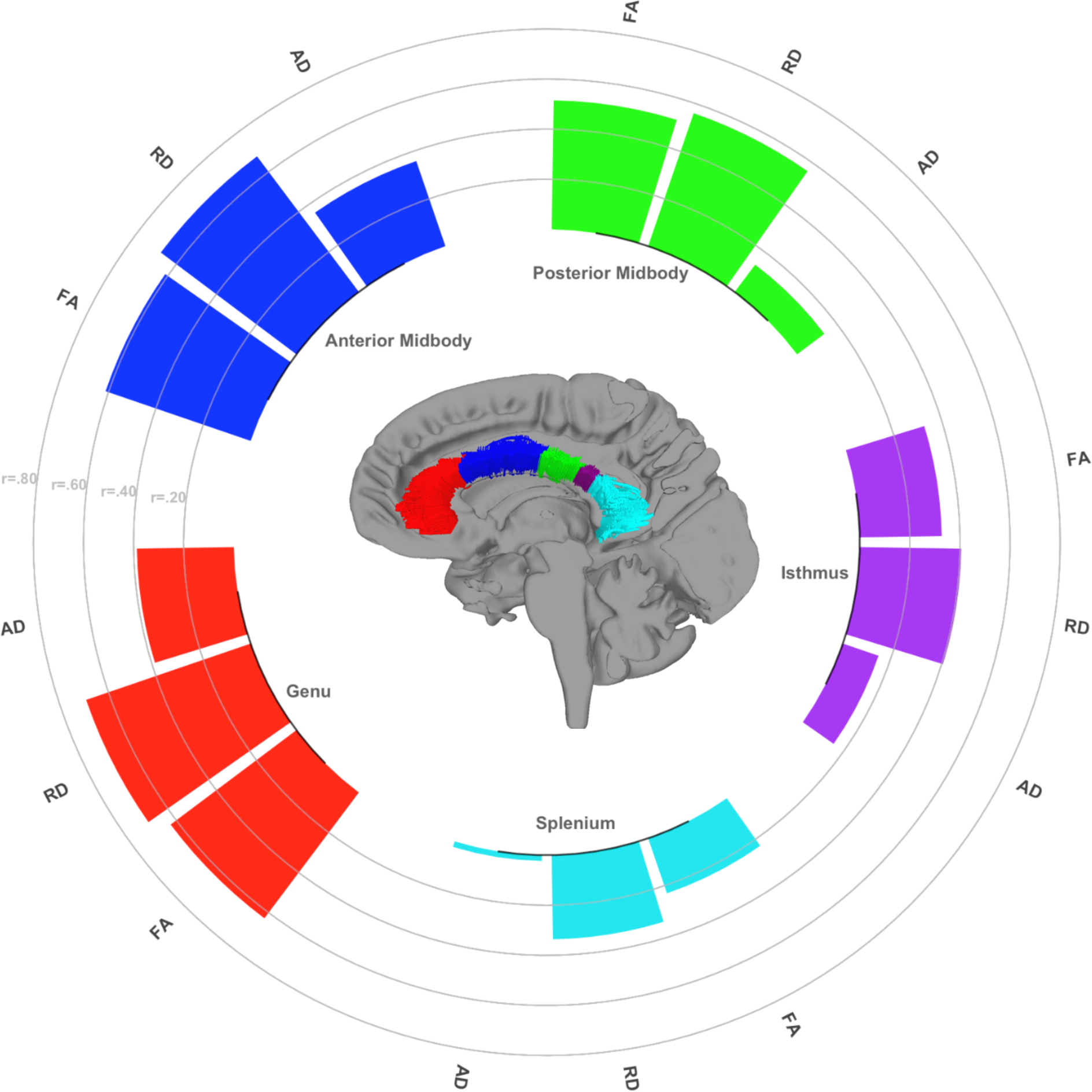
Circular barplot depicting gradation of the effects of age on FA, RD, and AD across CC subregion tracts. An anterior-to-posterior gradient of vulnerability to age is visible across the CC white matter. Steiger’s Z tests were used to compare significant differences between dependent correlations.

### Effects of Pulse Pressure on Regional CC Tract Metrics

We next performed GLMs using pulse pressure levels and age (centered at the mean as continuous variables) and their interaction as factors, and ROI as a within-subject dependent factor for each of the three diffusion metrics. In these models, we observed a main effect of age on FA (F(1,173)=36.67.121, p<0.0001) and RD (F(1,172)=44.03, p<0.0001), but not AD. There was a significant main effect of pulse pressure on RD (F(1,172)=4.98, p=0.027), with increasing pressure associated with increasing diffusivity. A significant age x pulse pressure interaction was observed for AD (F(2,171)=9.98, p=0.002). Within-subject, the age x ROI interaction observed previously in the age models remained significant for FA (F(1,172)=8.42, p<0.0001) and RD (F(1,171)=11.83, p<0.0001), but not AD (F(1,172)=1.93, p=0.11). As the effects of pulse pressure were not regionally selective, we used mean values of RD and AD across the entire corpus callosum for our secondary analyses. When age was parceled out of the association between PP and RD, the association remained significant (r_partial_(172)=0.17, p=0.026), see Figure 6.

**Figure 6.**
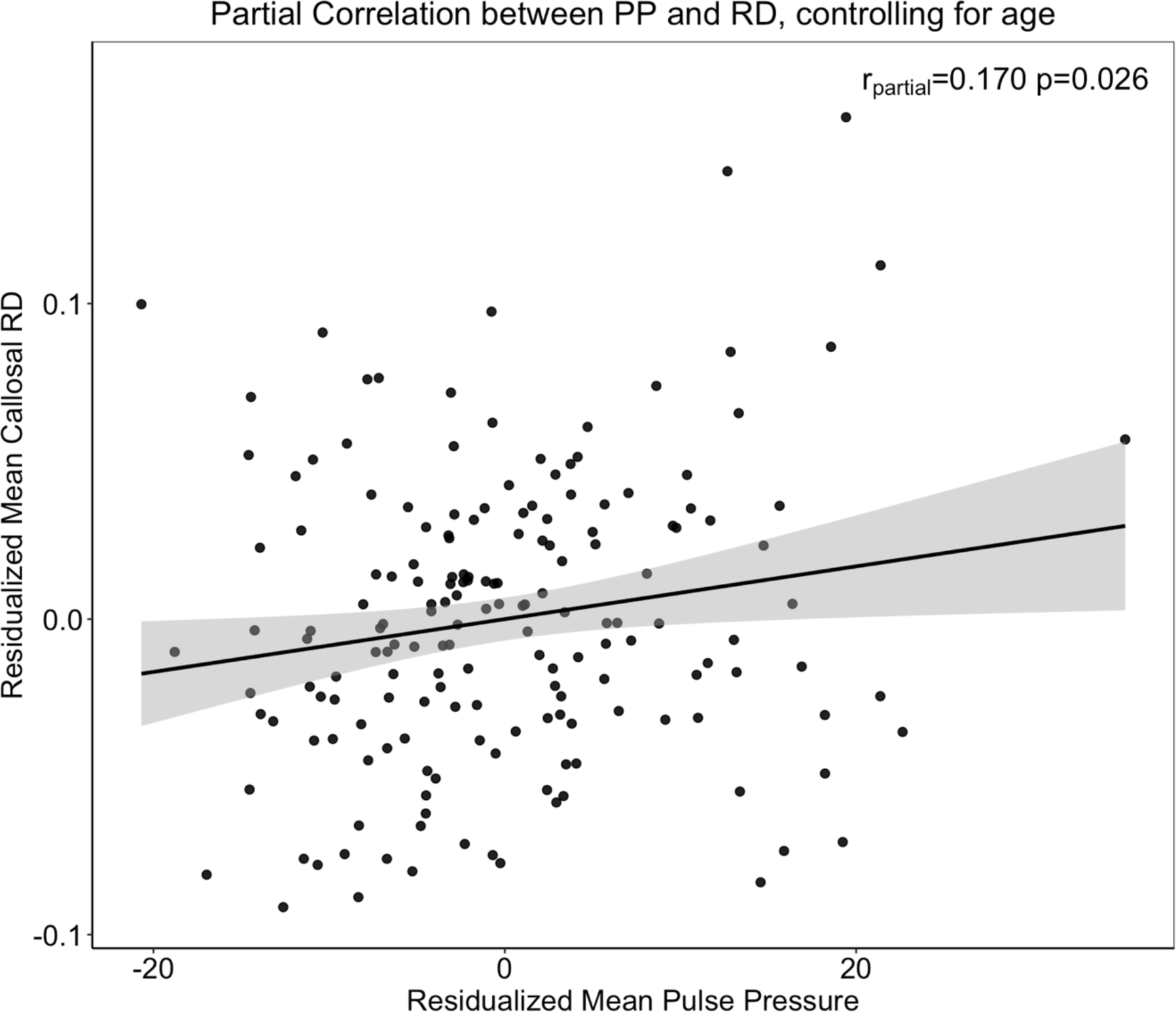
Significant association between pulse pressure and radial diffusivity, controlling for age. Radial diffusivity across the corpus callosum segments combined increases as a function of increasing pulse pressure, beyond the effects of age on RD.

To break down the interaction between age and pulse pressure on AD, we used the Johnson-Neyman method with simple slopes for visualization. Slopes were estimated at three levels: −1 standard deviation (SD) below the mean (34 years old), at the mean age (53 years old) and + 1 SD above the mean (54 years old) (Figure 7). Decomposition of the interaction reveals that the association between age and pulse pressure is primarily driven by the older adults (i.e., those +1 SD above the mean), r(172)=0.30, p=0.01, illustrated in Figure 7a. Johnson-Neyman interval calculation suggests that the age by pulse pressure interaction on AD is significant for participants beginning around the age of ∼60 years old (Figure 7b).

**Figure 7.**
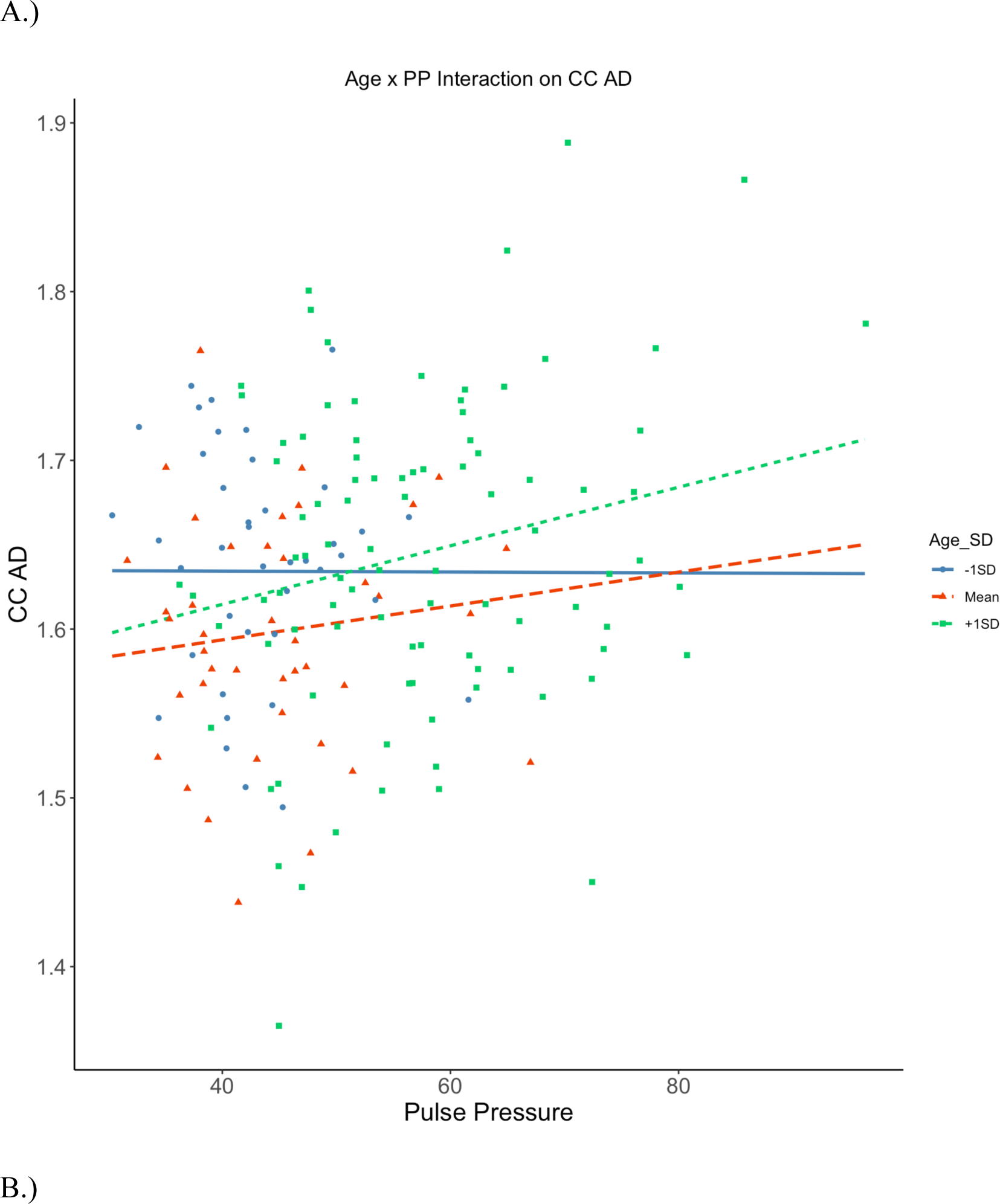
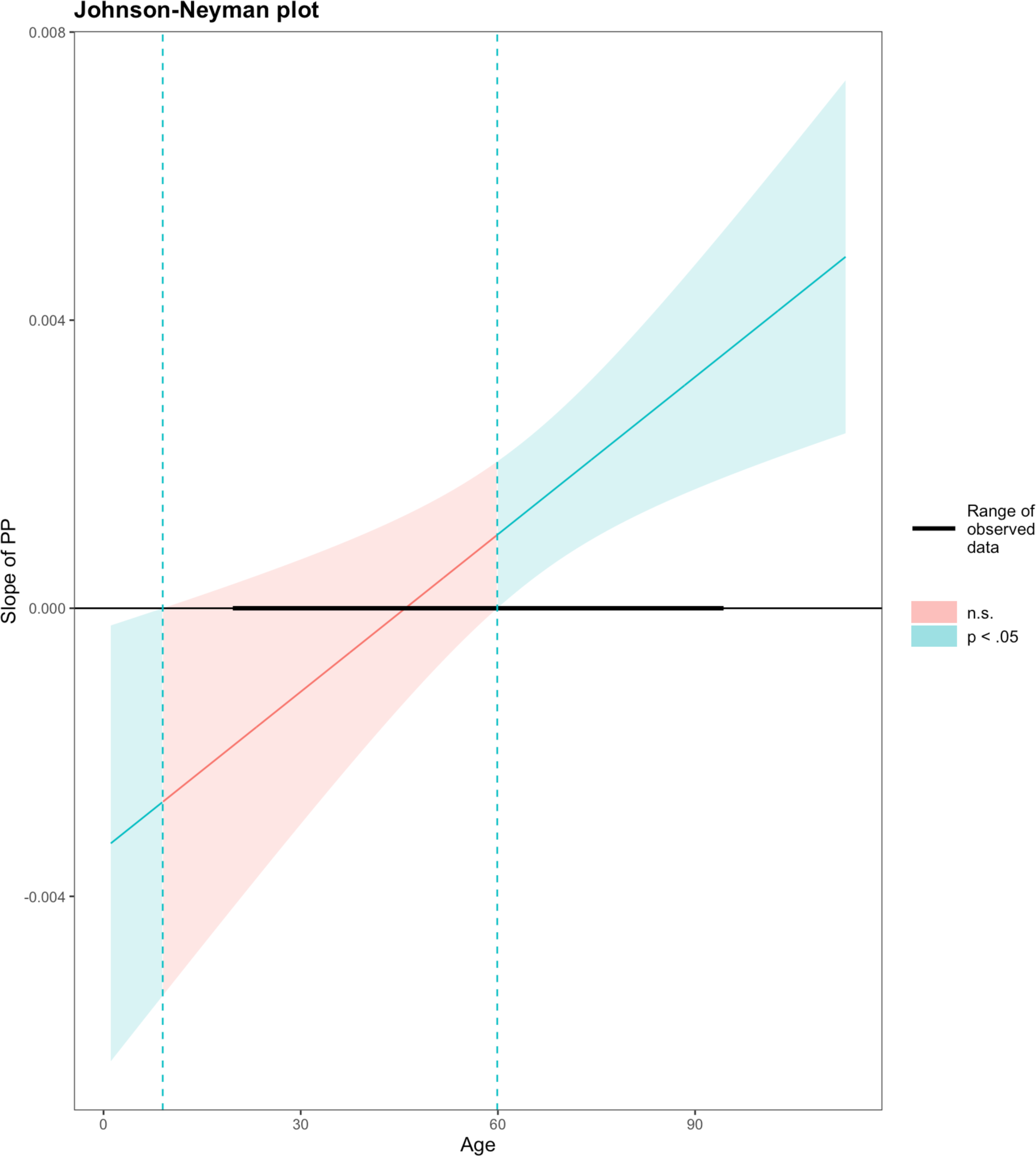
Age by Pulse Pressure Interaction on CC axial diffusivity. A). Simple slopes plots illustrating the effects of pulse pressure on AD are dependent on age, with significance in middle-aged and older adults, where increasing pulse pressure is associated with increased axial diffusivity in the corpus callosum. B). Johnson-Neyman intervals plot suggests that the significance range of the interaction begins at approximately the age of 60.

### Effects of Hypertension on Regional CC Tract Metrics

We found no significant effects of hypertensive status on FA, RD, or AD within the subregions of the CC (all p-values > 0.05).

## Discussion

In the present study, we replicate findings that the white matter fibers of the corpus callosum are highly vulnerable to the aging process. Moreover, our results demonstrate that the callosal subregions are differentially sensitive to age-related deterioration, as strength of the associations between age and FA uniformly decrease from genu to splenium, echoing the anterior-to-posterior gradient observed in white matter across the brain (Bennet et al., 2010; Davis et al., 2009; Kennedy & Raz, 2009b). According to the “last in, first out” hypothesis (Raz, 2000), this anterior-to-posterior gradient is a reversal of the pattern of white matter development, in which prefrontal regions are among the last to complete myelination. Associations between age and RD also followed the anterior-to-posterior gradient in our findings, and anterior ‘late-myelinating’ subregions had the greatest percentage of variance explained by age-related changes in RD. Overall, age was more strongly associated with RD than AD, suggesting that de/dysmyelination may be more integral to normal aging processes than total axonal degeneration, which may require frank injury or pathology to the brain. Other researchers have noted the opposite (Vernooij et al., 2008), and attributed white matter changes primarily to axonal degeneration and loss of macrostructural white matter organization. It may be that differing patterns of RD and AD alterations denote dissociable biological aging mechanisms (Bartzokis et al., 2012), and that these different processes are temporally and regionally specific (Burzynska et al., 2010). The non-linearity of age effects on both RD and AD in our findings has been noted in other studies (Hasan et al., 2009; Hsu et al., 2010). The linearity of age-related FA changes could be due to the exclusion of individuals under age 20, as Lebel et al. (2010) reported that FA values in the callosum peak during young adulthood.

The null finding for effects of hypertension as a diagnostic category on callosal white matter contradicts results from several previous studies that demonstrated an influence of hypertension diagnosis on both diffusivity and lesion load in the callosum (Burgman et al, 2010; Gons et al., 2015; Kennedy & Raz, 2009b). It could be that as a dichotomous diagnostic entity, hypertension is a more general factor compared to pulse pressure, which is the difference between systolic and diastolic blood pressure and is representative of arterial stiffness in the walls of larger arteries (Franklin, Khan, Wong, Larson, & Levy, 1999), and reflects a more specific, potentially mechanistic factor (arterial stiffness). In our study, we found that pulse pressure was associated with a negative, regionally invariant pattern of callosal microstructure. Specifically, pulse pressure appears to disrupt the anterior-to-posterior aging gradient, as differential (within-subject) effects of age were lost when accounting for pulse pressure. This may be attributable to de/dysmyelination in posterior subregions, as pulse pressure only significantly correlated in the isthmus and splenium for RD. Additionally, we found that the interaction between age and pulse pressure on AD was significant beginning around the age of 60 (Figure 7b). These findings suggest that pulse pressure begins to impact white matter health in the callosum by late midlife / early old age. The neural mechanisms underlying vascular health’s influence on white matter are not well understood, but the extant literature points to reduced cerebral blood perfusion and ischemic events as factors (Jefferson et al., 2011). Whatever mechanisms are responsible, they may represent a form of pathological aging, as poor vascular health is considered a major risk factor for vascular dementia (O’Brien and Thomas, 2015), and to a lesser degree Alzheimer’s Disease (Rodrigue et al., 2013).

A longer duration of hypertension diagnosis has been associated with greater age-related deterioration of posterior white matter areas, whereas elevations in pulse pressure, in a solely normotensive population, was linked to greater deterioration in anterior regions only (Kennedy and Raz (2009b), and replicated in other studies (Maillard et al., 2012; Salat et al., 2012). Additionally, after controlling for characteristics of small vessel disease (particularly white matter hyperintensity burden), at least one study (Gons, et al., 2012) reported that the association between hypertension and fractional anisotropy only remained significant in the splenium. Small increments in blood pressure levels may exacerbate age effects in the already compromised anterior subregions, and at higher levels, induce them in the relatively preserved posterior subregions, thereby disrupting the anterior-to-posterior gradient in typical aging.

The regional variability in callosal aging, and the modifying influence of vascular health, may have implications for certain cognitive functions that are thought to be dissociable for individual subregions. Executive functions associated with the prefrontal connections through the genu are among the cognitive abilities most sensitive to age; declines in genu FA, but not splenium FA, are related to poorer working memory (Kennedy & Raz, 2009a). In contrast, motor coordination is solely associated with FA in the body of the callosum (Johansen-Berg, Della-Maggiore, Behrens, Smith, & Paus, 2007), which connects premotor in the anterior body, and primary motor through the posterior midbody. These dissociations are due to the fact that mainly unimodal cortical projections pass through each subregion (Hofer and Frahm, 2006). In the presence of vascular risk factors, cognitive functions mediated by anterior regions may face even steeper declines, while cognitive domains relying on more posterior regions that are usually spared in normal aging may now also be compromised. Only limited evidence exists on the relationship between vascular health, white matter microstructure, and cognition (e.g., Li et al., 2015; Wang et al., 2015; Hoagey, Lazarus, Rodrigue & Kennedy, 2021), and future research should explore the impact of vascular health interventions on susceptibility to callosal microstructure degradation across the lifespan. For example, one way that exercise positively affects brain health and cognition is by increasing cerebral blood flow (e.g., Tarumi et al., 2022; Tomoto et al., 2023).

There are several limitations to the study that must be considered. First, this is a cross-sectional report, and further longitudinal investigations are needed to verify whether the patterns of aging present in our findings are representative of true within-person changes in aging. Such data are currently being collected in follow-up waves to this study. Second, as this study was not designed to specifically study hypertension, there are some methodological considerations. We were unable to control or test for the effects of antihypertensive medication use, as we did not have a large enough sample of diagnosed hypertensives to sufficiently divide into separate medication subsamples. Third, diffusion tensor imaging has its own limitations (e.g., Jones & Cercignani, 2010; Figley et al., 2022), although careful quality checks, visual inspection of all images, and proper tractography methods utilized in this study were intended to mitigate those limitations. Many of the limitations from diffusion tensor models are most notable in complex white matter areas of intermixing fiber types, and less applicable to the fibers comprising the corpus callosum.

In conclusion, aging differentially and negatively impacts the microstructural health of the subregions of the corpus callosum, and elevations in pulse pressure may alter these patterns of aging, even in individuals without diagnosed hypertension. Maintaining optimal blood pressure levels to avoid or delay age-related arterial stiffening could be a key factor to successful aging, but more research is needed in this area to determine how vascular health interacts with age to influence brain structure and cognition.

## Notes

### Competing Interest Statement

The authors have declared no competing interest.

